# The Lipid Elongation Enzyme ELOVL2 is a molecular regulator of aging in the retina

**DOI:** 10.1101/795559

**Authors:** Daniel Chen, Daniel L. Chao, Lorena Rocha, Matthew Kolar, Viet Anh Nguyen Huu, Michal Krawczyk, Manish Dasyani, Tina Wang, Maryam Jafari, Mary Jabari, Kevin D. Ross, Alan Saghatelian, Bruce Hamilton, Kang Zhang, Dorota Skowronska-Krawczyk

**Author notes:** these authors contributed equally (Daniel Chen, Daniel L. Chao).

## Abstract

Methylation of the regulatory region of the Elongation Of Very Long Chain Fatty Acids-Like 2 (*ELOVL2*) gene, an enzyme involved in elongation of long-chain polyunsaturated fatty acids, is one of the most robust biomarkers of human age, but the critical question of whether *ELOVL2* plays a functional role in molecular aging has not been resolved. Here, we report that *Elovl2* regulates age-associated functional and anatomical aging *in vivo*, focusing on mouse retina, with direct relevance to age-related eye diseases. We show that an age-related decrease in *Elovl2* expression is associated with increased DNA methylation of its promoter. Reversal of *Elovl2* promoter hypermethylation *in vivo* through intravitreal injection of 5-Aza-2’-deoxycytidine (5-aza-dc) leads to increased *Elovl2* expression and rescue of age-related decline in visual function. Mice carrying a point mutation C234W that disrupts *Elovl2*-specific enzymatic activity show electrophysiological characteristics of premature visual decline, as well as early appearance of autofluorescent deposits, well-established markers of aging in the mouse retina. Finally, we find deposits underneath the retinal pigment epithelium in *Elovl2* mutant mice, containing components of complement system and lipid metabolism. These findings indicate that ELOVL2 activity regulates aging in mouse retina, provide a molecular link between polyunsaturated fatty acids elongation and visual functions, and suggest novel therapeutic strategies for treatment of age-related eye diseases.

## INTRODUCTION

Chronological age predicts relative levels of mental and physical performance, disease risks across common disorders, and mortality ^1^. The use of chronological age is limited, however, in explaining the considerable biological variation among individuals of a similar age. Biological age is a concept that attempts to quantify different aging states influenced by genetics and a variety of environmental factors. While epidemiological studies have succeeded in providing quantitative assessments of the impact of discrete factors on human longevity, advances in molecular biology now offer the ability to look beyond population-level effects and to hone in on the effects of specific factors on aging within single organisms.

A quantitative model for aging based on genome-wide DNA methylation patterns by using measurements at 470,000 CpG markers from whole blood samples of a large cohort of human individuals spanning a wide age range has recently been developed ^2–4^. This method is highly accurate at predicting age, and can also discriminate relevant factors in aging, including gender, genetic variants, and disease ^2,5^. Several models work in multiple tissues ^3,4^, suggesting the possibility of a common molecular clock, regulated in part by changes in the methylome. In addition, these methylation patterns are strongly correlated with cellular senescence and aging ^6^. The regulatory regions of several genes become progressively methylated with increasing chronological age, suggesting a functional link between age, DNA methylation, and gene expression. The promoter region of *ELOVL2*, in particular, was the first to be shown to reliably show increased methylation as humans age ^7^, and confirmed in the one of the molecular clock models ^2^.

*ELOVL2* (Elongation Of Very Long Chain Fatty Acids-Like 2) encodes a transmembrane protein involved in the elongation of long-chain (C22 and C24) omega-3 and omega-6 polyunsaturated fatty acids (LC-PUFAs)^8^. Specifically, ELOVL2 is capable of converting docosapentaenoic acid (DPA) (22:5n-3) to 24:5n-3, which can lead to the formation of very long chain PUFAs (VLC-PUFAs) as well as 22:6n-3, docosahexaenoic acid (DHA) ^9^. DHA is the main polyunsaturated fatty acid in the retina and brain. Its presence in photoreceptors promotes healthy retinal function and protects against damage from bright light and oxidative stress. *ELOVL2* has been shown to regulate levels of DHA ^10^, which in turn has been associated with age-related macular degeneration (AMD), among a host of other retinal degenerative diseases ^11^. In general, LC-PUFAs are involved in crucial biological functions including energy production, modulation of inflammation, and maintenance of cell membrane integrity. It is, therefore, possible that *ELOVL2* methylation plays a role in the aging process through the regulation of these diverse biological pathways.

In this study, we investigated the role of ELOVL2 in molecular aging in the retina. We find that the *Elovl2* promoter region is increasingly methylated with age in the retina, resulting in age-related decreases in *Elovl2* expression. These changes are associated with decreasing visual structure and function in aged mice. We then demonstrate that loss of ELOVL2-specific function results in the early-onset appearance of sub-RPE deposits that contain molecular markers found in drusen in AMD. This phenotype is also associated with visual dysfunction as measured by electroretinography, and it suggests that ELOVL2 may serve as a critical regulator of a molecular aging clock in the retina, which may have important therapeutic implications for diseases such as age-related macular degeneration.

## RESULTS

### Elovl2 expression is downregulated with age through methylation and is correlated with functional and anatomical biomarkers in aged wildtype mice

Previous studies showed that methylation of the promoter region of *ELOVL2* is highly correlated with human age ^2^. Methylation of regulatory regions is thought to prevent the transcription of neighboring genes and serves as a method to regulate gene expression. We first wished to characterize whether the age-associated methylation of the *ELOVL2* promoter previously found in human serum also occurs in the mouse. First, we analyzed ELOVL2 promoter methylation data obtained using bisulfite-sequencing in mouse blood and compared it to the available human data for the same region ^12^ and observed similar age-related increase in methylation level in the compared regions (**Figure S1A**). To assay methylation of the *Elovl2* promoter in retina, we used methylated DNA immunoprecipitation (MeDIP) method ^13^ and tested the methylation levels in the CpG island in the *Elovl2* regulatory region by quantitative PCR with *Elovl2*-specific primers (**Supp. Table 1**). MeDIP analysis of the CpG island in the *Elovl2* regulatory region showed increasing methylation with age in the mouse retina (**Fig. 1A**). This was well-correlated with age-related decreases in expression of *Elovl2* as assessed by Western blot and qPCR (**Fig. 1B and Fig S1B,C**) indicating the potential role of age-related changes in DNA methylation in *Elovl2* expression.

**Figure 1.**
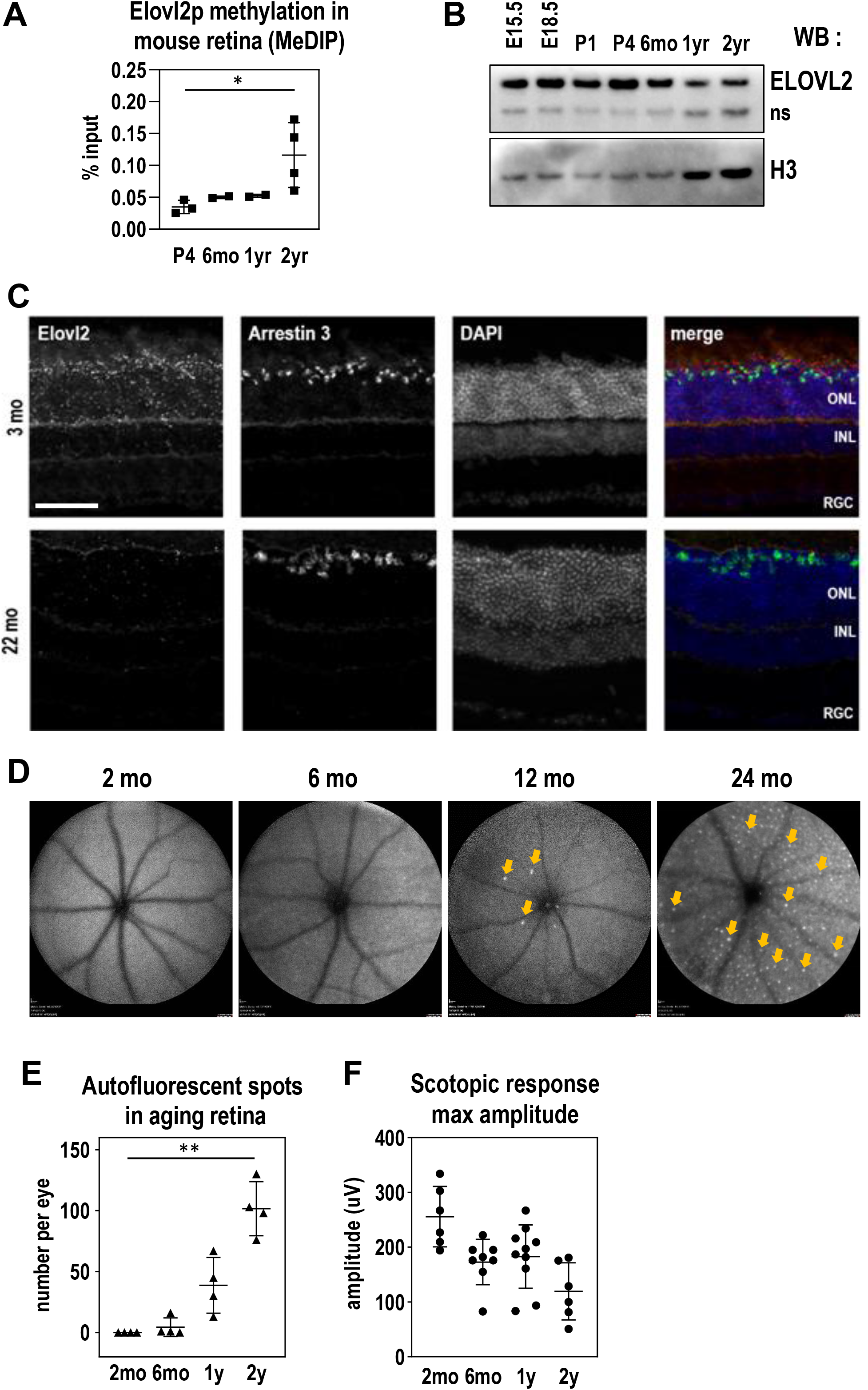
ELOVL2 expression is downregulated with age through methylation of its promoter and is correlated with age related increases in autofluorescence aggregates and decreased scotopic response.

**A.** Methylation of ELOVL2 promoter region measured using immunoprecipitation of methylated (MeDIP) followed by qPCR. ELOVL2 promoter is increasingly methylated with age. **B**. Time course of retinal ELOVL2 protein expression by Western blot. ELOVL2 protein is expression decreases with age. ns, non-specific signal produced by ELOVL2 antibodies **C**. Images of mouse retina sections from young – 3mo (top panels) and old – 22mo (bottom panels) animals stained with RNA-scope probes designed for *Elovl2* and *Arrestin 3*, counterstained with DAPI. ONL, outer nuclear layer, INL, inner nuclear layer, RGC, retinal ganglion cells. Bar −100um **D**. Time course of representative fundus autofluorescence pictures of C57BL/6J mice. Arrows denote autofluorescent deposits. **E**. Quantification of autofluorescent deposits in fundus images. N=4. **F**. Scotopic responses by ERG over mouse lifespan. For panels A, E and F: N=4, *p<0.5, ** p <0.01, 1-way ANOVA. Error bars denote SD.

To understand the cell-type and age-specific expression of *Elovl2*, we performed *in situ* hybridization with an *Elovl2* RNAscope probe on mouse retina sections ^14^. In three-month-old and in 22-month-old mice, we noticed *Elovl2* expression in the photoreceptor layer, particularly in the cone layer as well as the RPE (**Fig. 1C and Fig. S1E**). We observed that the expression of Elovl2 on mRNA level in RPE was lower than in the retina (**Fig. S1D**). Importantly, at older stages (22-month animals), we noticed *Elovl2* mRNA in the same locations but dramatically reduced in expression (**Fig. 1C**). As *Elovl2* is also highly expressed in the liver, we performed a time course of *Elovl2* expression in this tissue. We observed similar age-related decreases in *Elovl2* expression correlated with increases in methylation of the *Elovl2* promoter in mouse liver, indicating that age-associated methylation of *Elovl2* occurs in multiple tissues in mice (**Fig. S1F**).

Visual function is highly correlated with age, including age-related decreases in rod function in both humans and mice ^15,16^. In addition, autofluorescent aggregates have been observed in the fundus of aged mice, suggesting that these aggregates may also be an anatomical surrogate of aging in the mouse retina ^17,18^. To measure and correlate these structural and visual function changes with age in mice, we performed an analysis of wildtype C57BL/6J mice at various timepoints through development, using fundus autofluorescence and electroretinography (ERG) as structural and functional readouts for vision. We observed increasing amounts of autofluorescent aggregates on fundus autofluorescence imaging with increasing mouse age, most prominently at two years (**Fig. 1D, E and Fig. S1G**). We also detected an age-associated decrease in visual function, as measured by maximum scotopic amplitude by ERG (**Fig. 1F and Fig. S1H**), as shown in previous studies ^15,19^. These data show that an age-associated accumulation of autofluorescent spots and decrease of visual function as detected by ERG correlate with *Elovl2* downregulation in the mouse retina.

### Manipulating ELOVL2 expression causes age-related changes in cells

The WI38 and IMR90 cell lines are well-established cell models of aging ^20^. We used these cell lines to further explore the effect of *ELOVL2* promoter methylation on cell health. First, using MeDIP, we found that promoter methylation increased with cell population doubling (**Fig. 2A**) further confirming strong correlation between increased *ELOVL2* methylation and aging. Since the methylation of the promoter region was shown to be inhibitory for transcription ^21^, we investigated whether the expression level of *ELOVL2* inversely correlated with *ELOVL2* promoter methylation. Using qRT-PCR, we found that the expression level of the gene decreased with increasing population doubling (PD) number (**Fig. 2B**). We conclude that *ELOVL2* expression is downregulated in aging cells, with a correlated increase in *ELOVL2* promoter methylation.

**Figure 2.**
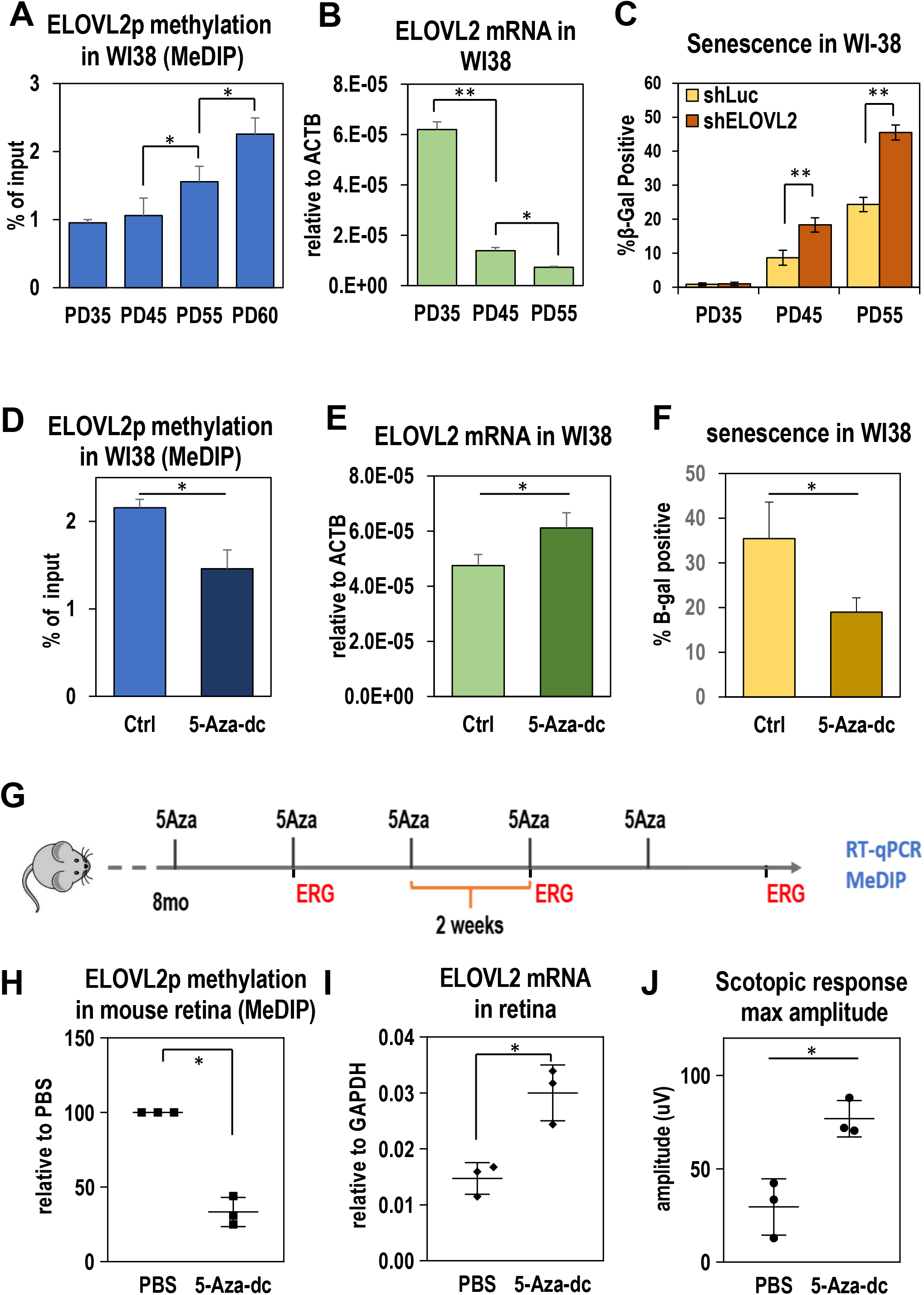
A-C, *ELOVL2* expression, methylation and senescence in WI38 cells.

**A.** Methylation level in *ELOVL2* promoter region in human normal lung cell line WI38 by MeDIP/qPCR. Amplicons contain CpG markers cg16867657, cg24724428, and cg21572722. N>3 **B**. *ELOVL2* expression by qPCR in WI38 cells at PD35, 45, 55. **C**. Fraction of senescent cells measured by beta-galactosidase staining in WI38 cells at given population doubling upon shRNA mediated knock-down of ELOVL2 gene or control Luc. **D-F,** Manipulating DNA methylation in PD52 WI38 cells. **D**. *ELOVL2* promoter methylation as measured by MeDIP followed by qPCR in untreated control and 5-Aza-dc treated WI38 cells. **E**. *ELOVL2* expression by qPCR in untreated control and 5-Aza-dc treated WI38 cells. **F**. Percent senescence by beta-galactosidase staining in WI38 cells treated with 2μM 5-Aza-dc. **G-J**, Manipulating DNA methylation in mice. **G**. Experimental setup. 8 month old mice were injected intravitreally with of 5-Aza-dc five times every two weeks. ERG measurements were taken at indicated time points. At 11 months, expression and methylation levels were measured in 5-Aza-dc treated and control (PBS-treated) mice. **H**. Methylation of ELOVL2 promoter by MeDIP at 11 months after 5-Aza injection. **I**. *ELOVL2* expression by qPCR after 5-Aza injection. **J**. Maximum amplitude scotopic response by ERG after 5-Aza injection. For panels A-F N>=3, *p<0.05, **p<0.01, t-test. Error bars denote SD; for panels H-J N=3, *p<0.05, **p<0.01, t-test. Error bars denote SD.

We then asked whether modulating the expression of *ELOVL2* could influence cellular aging. First, using shRNA delivered by lentivirus, we knocked down *ELOVL2* expression in WI38 and another model cell line, IMR-90, and observed a significant decrease in proliferation rate (**Fig. S2A, B**), an increased number of senescent cells in culture as detected by SA-β-Gal staining (**Fig 2C and Fig. S2E**), and morphological changes consistent with morphology of high PD cells (**Fig. S2F**). Altogether, these data suggest that decreasing ELOVL2 expression results in increased aging and senescence *in vitro*.

Next, we tested whether we could manipulate *Elovl2* expression by manipulating the *Elovl2* promoter methylation. We treated WI38 fibroblasts with 5-Aza-2’-deoxycytidine (5-Aza-dc), a cytidine analog that inhibits DNA methyltransferase ^22^. Cells were treated for two days with 2 μM 5-Aza-dc followed by a five-day wash-out period. Interestingly, we found that upon treatment with 5-Aza-dc, *Elovl2* promoter methylation was reduced (**Fig. 2D**), and *Elovl2* expression was upregulated (**Fig. 2E**). Moreover, upon 5-Aza-dc treatment, a lower percentage of senescent cells were observed in culture (**Fig. 2F**). To assess whether the decrease of senescence is caused at least in part by the ELOVL2 function we knocked down the *ELOVL2* expression in aged WI38 cells and treated them with 5-Aza-dc as previously described. Again, significantly lower proportion of senescent cells was detected upon the drug treatment, but the effect of drug treatment was significantly reduced by shRNA-mediated knockdown of *ELOVL2*, using either of two *ELOVL2* shRNAs compared to a control shRNA (**Figure S2B**). This indicates an important role of ELOVL2 in the process. Altogether, these data suggest that the reversing *ELOVL2* promoter methylation increases its expression and decreases senescence *in vitro*.

### DNA demethylation in the retina by intravitreal injection of 5-Aza-dc increases Elovl2 expression and rescues age-related changes in scotopic function in aged mice

We next explored whether demethylation of the *Elovl2* promoter could have similar effects on *Elovl2* expression *in vivo*. To accomplish this, we performed intravitreal injection of 5-Aza-dc, known to affect DNA methylation in nondividing neurons^23–25 26^, into aged wildtype mice. 8-month-old C57BL/6J mice were injected with 1 μL of 2 μM 5-Aza-dc in one eye and 1 μL of PBS in the other eye as a control, every other week over a period of 3 months (total of 5 injections) (**Fig. 2G**). After the treatment, tissues were collected, and RNA and DNA were extracted. We found, using the MeDIP method, that methylation of the *Elovl2* promoter decreased after treatment (**Fig. 2H**), with a corresponding upregulation of *Elovl2* expression (**Fig. 2I**). Notably, we observed that the scotopic response was significantly improved in the 5-Aza-dc injected eyes compared to vehicle controls (**Fig. 2J**). These data show that DNA demethylation, which included demethylation of the *Elovl2* promoter region, influence and potentially delay age-related changes in visual function in the mouse retina.

### Elovl2^C234W^ mice demonstrate a loss of ELOVL2-specific enzymatic activity

We next sought to investigate the *in vivo* function of *Elovl2* in the retina. Since C57BL/6 *Elovl2* +/-mice display defects in spermatogenesis and are infertile^27^, we developed an alternative strategy to eliminate ELOVL2 enzymatic activity *in vivo*. Using CRISPR-Cas9 technology, we generated *Elovl2*-mutant mice encoding a cysteine-to-tryptophan substitution (C234W). This mutation selectively inactivates enzymatic activity of *ELOVL2* required to process C22 PUFAs, to convert docosapentaenoic acid (DPA) (22:5n-3) to 24:5n-3, while retaining elongase activity for other substrates common for ELOVL2 and the paralogous enzyme ELOVL5 (**Fig. 3A, Fig. S3A**)^9,28,29^. A single guide RNA against the *Elovl2* target region, a repair oligonucleotide with a base pair mutation to generate the mutant C234W, and Cas9 mRNA were injected into C57BL/6N mouse zygotes (**Fig. 3B**). One correctly targeted heterozygous founder with the C234W mutation was identified. No off-target mutations were found based on DNA sequencing of multiple related DNA sequences in the genome. (**Fig. S3B**). The C234W heterozygous mice were fertile, and C234W homozygous mice developed normally and showed no noticeable phenotypes. We analyzed the long chain fatty levels in the retinas of homozygous *Elovl2*^C234W^ mice to determine whether there was a loss of enzymatic activity specific to ELOVL2. We observed that *Elovl2*^C234W^ mice had higher concentrations of C22:5 fatty acid (a selective substrate of ELOVL2 elongation) and lower levels of C24:5 (primary product of ELOVL2 enzymatic activity) and C22:6 (DHA – the secondary product of ELOVL2) (**Fig. 3C**). We also observed similar changes in fatty acid levels in livers of *Elovl2*^C234W^ mice as well as lower levels of longer fatty acids that require primary product of Elovl2 as a substrate (**Fig. S4**) This suggests that the *Elovl2*^C234W^ mice have altered ELOVL2 substrate specificity and inhibited ELOVL2-specific C22 elongase activity.

**Figure 3.**
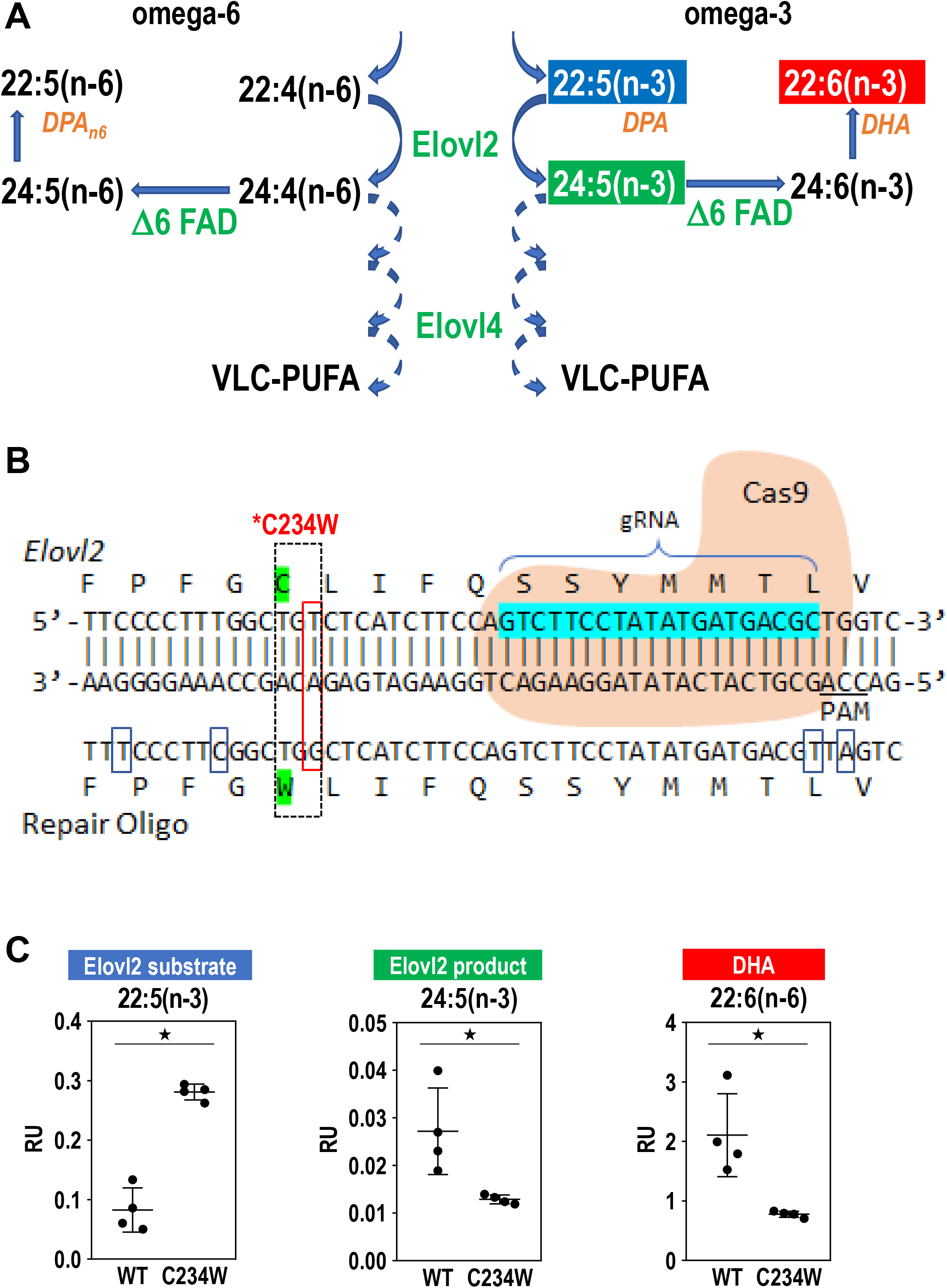
Elovl2^C234W^ mice show a loss of ELOVL2 enzymatic activity.

**A.** Schematic of ELOVL2 elongation of omega-3 and omega-6 fatty acids. ELOVL2 substrates 22:5 (n-3) and 22:4(n-6) are elongated by ELOVL2 to 24:5(n-3) and 24:4(n-6). This leads to other products such as DHA, DPAn6 as well as VLC-PUFAs, which are elongated by ELOVL4. **B**. CRISPR-Cas9 strategy to create *Elovl2*^C234W^ mice. *Elovl2* gRNA, Cas9 and repair oligo are used to create the *Elovl2*^C234W^ mutant. **C**. Lipid levels of ELOVL2 substrate DPA (22:5(n-3)), ELOVL2 product (24:5(n-3)), and DHA (22:6(n-3)) in retinas of *Elovl2*^C234W^ mice and wild-type littermates. N=4, *p<0.05 by Mann-Whitney U-test. Error bars represent SD.

### Loss of ELOVL2-specific activity results in early vision loss and accumulation of subRPE deposits

We next investigated whether the *Elovl2*^C234W^ mutation affected the retinal structure and/or function *in vivo*. First, we observed a significant number of autofluorescent spots on fundus photography in animals at six months of age, which were not found in wild-type littermates (**Fig 4A, B**). This phenotype was consistently observed in 6, 8, and 12-month old mutant animals and in both animal sexes, but the phenotype was consistently more pronounced in male mice (**Fig. S5**). Importantly, ERG analysis revealed that 6-month old *Elovl2*^C234W^ mice displayed a decrease in visual function as compared to wild type littermates (**Fig. 4C, Fig. S5**).

**Figure 4.**
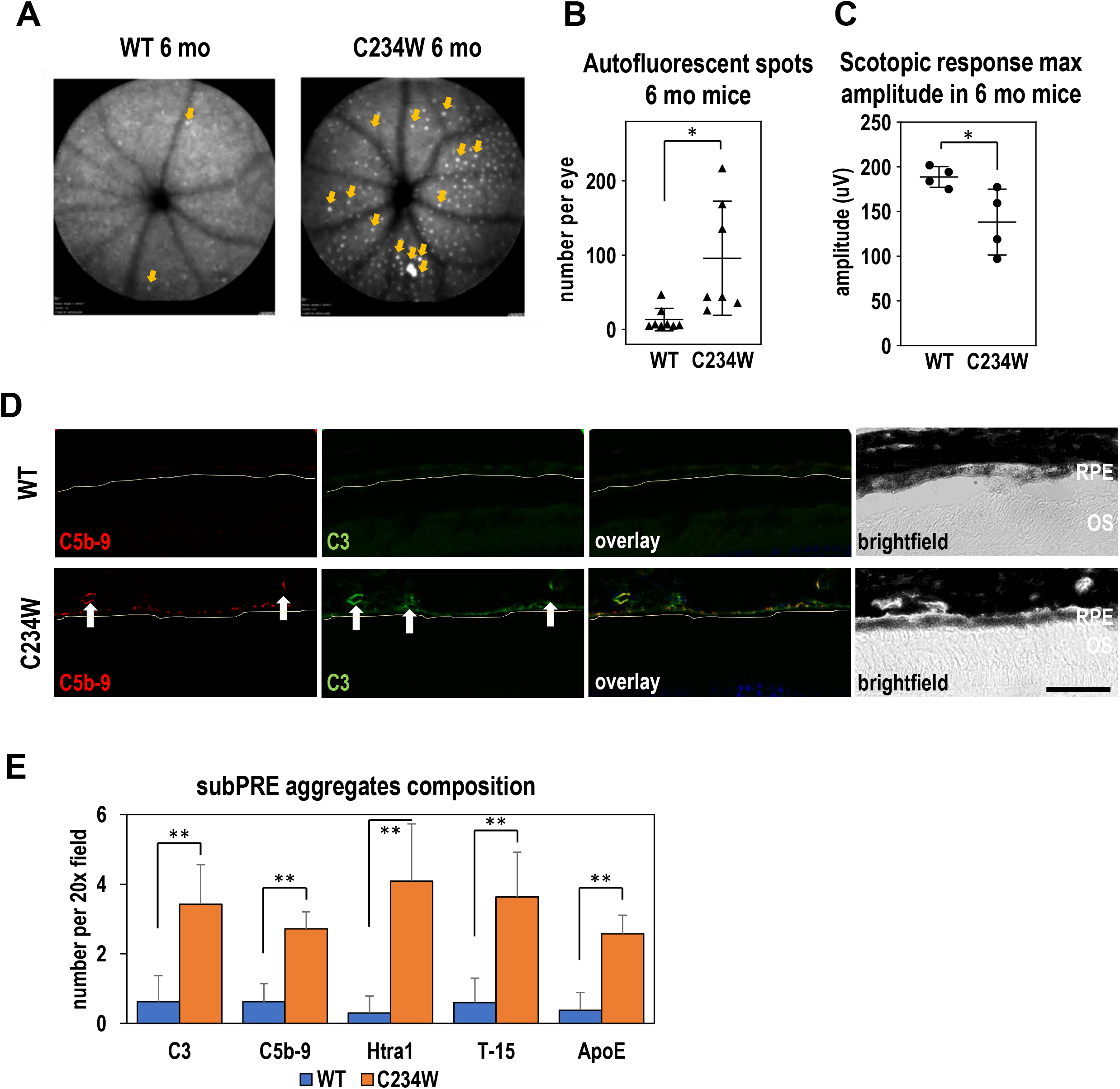
*Elovl2*^C234W^ mice show autofluorescent deposits and vision loss.

**A**. Representative fundus autofluorescence images of WT and *Elovl2*^C234W^ mice at 6 months with representative scotopic ERG waveforms. Note multiple autofluorescent deposits (arrows) in *Elovl2*^C234W^ mice which are almost absent in wild-type littermates. **B**. Quantification of the autofluorescent spots in 6mo wild-type and C234W mutant mice. N=8. *p<0.05, t-test. Error bars denote SD. **C**. Maximum scotopic amplitude by ERG at 6 months between WT and *Elovl2*^C234W^ mice. N=4, *p<0.05, t-test. Error bars represent SD. **D**. Immunohistochemistry of sub-RPE deposits found in *Elovl2*^C234W^ mice. Deposits are found underneath the RPE (yellow line), which colocalize with C3 and C5b-9, which is not present in WT controls. Bar – 50um. **E**. Quantification of subRPE aggregates stained with C3, C5b-9, Htra1, T-15, and ApoE, all components found in drusen in AMD. N=4, ** p<0.01, t-test. Error bars represent SD.

To determine the impact of the mutation on the morphology of the retina on the microscopic level, we performed an immunohistological analysis of tissue isolated form wild type and *Elovl2*^C234W^ littermates. Although we did not observe gross changes in morphology of the retinas in mutant animals, we have observed the presence of small aggregates underneath the RPE and found that these subRPE aggregates contained the complement component C3 as well as the C5b-9 membrane attack complex, proteins found in human drusenoid aggregates (**Figure 4D**). In addition, in the mutant subRPE aggregates, we also identified other components found in human deposits such as HTRA1^30^, oxidized lipids T-15^31^, and ApoE, an apolipoprotein component of drusen^32^ (**Figure 4E**). This suggests that the subRPE deposits found in the *Elovl2*^C234W^ mouse contain some drusen-specific components found in early nonexudative AMD. Taken together, these data implicate ELOVL2-specific activity as a potential functional target in age-related eye diseases.

## DISCUSSION

### ELOVL2 as a critical regulator of molecular aging in the retina

This work is the first demonstration, to our knowledge, of a functional role for *Elovl2* in regulating age-associated phenotypes in the retina. Methylation of the promoter region of *ELOVL2* is well established as a robust prognostic biomarker of human aging^7,33^, but whether ELOVL2 activity contributes to aging phenotypes had not yet been documented. In this work, we demonstrated that the age-related methylation of regulatory regions of *Elovl2* occurs in the rodent retina and results in age-related decreases in the expression of *Elovl2*. We show that inhibition of *ELOVL2* expression by transfection of *ELOVL2* shRNA in two widely-used cell models results in increased senescence and decreased proliferation, endpoints associated with aging. Conversely, we show that the administration of 5-Aza-dc leads to demethylation of *ELOVL2* promoter and prevents cell proliferation and senescence compared to controls.

Next, we explored whether *Elovl2* expression affected age-related phenotypes *in vivo*. Intravitreal injection of 5-Aza-dc in rodents increased *Elovl2* expression and reversed age-related changes in visual function by ERG. Next, we showed a decrease in visual function as assessed by ERG as well as increased accumulation of autofluorescent white spots in *Elovl2^C234W^* mice, with ELOVL2-specific activity eliminated, compared to littermates controls. These physiologic and anatomic phenotypes are well-established markers of aging in the mouse retina, suggesting that loss of *Elovl2* may be accelerating aging on a molecular level in the retina. Finally, in *Elovl2*^C234W^ mice, we observed the appearance of sub-RPE deposits, which colocalize with markers found in human drusen in macular degeneration, a pathologic hallmark of a prevalent age-related disease in the eye. Taken together, we propose that *Elovl2* plays a critical role in regulating a molecular aging in the retina, which may have therapeutic implications for age-related eye diseases.

### Methylation of the regulatory region as a mechanism of age-dependent gene expression

DNA methylation at the 5-position of cytosine (5-methylcytosine, 5mC) is catalyzed and maintained by a family of DNA methyltransferases (DNMTs) in eukaryotes ^34^ and constitutes ~2-6% of the total cytosines in human genomic DNA (28). Alterations of 5mC patterns within CpG dinucleotides within regulatory regions are associated with changes in gene expression ^21,35^. Recently it has been shown that one can predict human aging using DNA methylation patterns. In particular, increased DNA methylation within the CpG island overlapping with the promoter of *ELOVL2* was tightly correlated with the age of the individual^33^. We attempted to demethylate this region using 5-Aza-dc, known to inhibit the function of DNMTs also in nondividing neurons^23–25^. We reported that upon intravitreal injection of the compound, the DNA methylation is reduced, gene expression is upregulated, and visual function is maintained in the treated eye compared with the contralateral control. These data suggest that *Elovl2* is actively methylated by enzymes inhibited by 5Aza-dc and that age-related methylation either directly or indirectly regulates *Elovl2* expression. Further studies are needed to fully address the directness and specificity of methylation effects on *Elolv2* expression and visual function.

### A molecular link between long-chain PUFAs in age-related eye diseases

Our data show that *Elovl2*^C234W^ animals display accelerated loss of vision and the appearance of macroscopic autofluorescent spots in fundus images. The exact identity of such spots in mouse models of human diseases is unclear, as they have been suggested to be either protein-rich, lipofuscin deposits or accumulating microglia^17,36^. Rather than deciphering the identity of these macroscopic spots, we used the phenotype as a potential sign of age-related changes in the retina, as suggested by others^17,37^.

The composition of aggregates visible on the microscopic level in sub-RPE layers in the retina is potentially informative with regard to human parallels. Using immunofluorescence we observed the accumulation of several proteins described previously as characteristic for drusen in human AMD samples. Although, our analysis did not exhaust the documented components of drusen in human disease^38^, nevertheless, our data show the appearance of these subRPE deposits, even in the absence of known confounding mutations or variants correlating with the risk of the disease.

What may be the mechanism by which *Elovl2* activity results in drusen-like deposits and loss of visual function? ELOVL2 plays an essential role in the elongation of long-chain (C22 and C24) omega-3 and omega-6 polyunsaturated acids (LC-PUFAs) (**Fig. 3A**). LC-PUFAs are found primarily in the rod outer segments and play essential roles in retinal function. These PUFAs include both long chain omega-3 (n-3) and omega-6 (n-6) fatty acids such as docosahexaenoic acid (DHA) and arachidonic acid (AA). DHA is the major polyunsaturated fatty acid found in the retina and has been shown to play diverse roles in photoreceptor function, protection in oxidative stress, as well as retinal development ^39^. While DHA has been well studied in the human retina, the function of other LC-PUFAs in the ELOVL2 elongation pathway is unknown. Further experiments to dissect the roles of specific LC-PUFAs in this pathway, and which of these lipid species are implicated in this phenotype are still required.

Multiple lines of evidence have linked PUFAs to age-related macular degeneration (AMD). AMD is the leading cause of blindness in developed countries^40^ among the elderly. There are two advanced subtypes of AMD, an exudative form due to neovascularization of the choroidal blood vessels, and a nonexudative form which results in gradual retinal pigment epithelium (RPE) atrophy and photoreceptor death. While there are currently effective therapies for exudative AMD, there are no treatments which prevent photoreceptor death from nonexudative AMD. A pathologic hallmark of non-exudative AMD is the presence of drusen, lipid deposits found below the RPE, which leads to RPE atrophy and photoreceptor death, termed geographic atrophy. The pathogenesis of macular degeneration is complex and with multiple pathways implicated including complement activation, lipid dysregulation, oxidative stress, and inflammation among others^40^. Despite intense research, the age-related molecular mechanisms underlying drusen formation and geographic atrophy are still poorly understood.

Analysis of AMD donor eyes showed decreased levels of multiple LC-PUFAs and VLC-PUFAs in the retina and RPE/choroid compared to age-matched controls^41^. Epidemiologic studies suggest that low dietary intake of LC-PUFAs such as omega-3 fatty acids was associated with a higher risk of AMD^42,43^. Furthermore, mutations in ELOVL4, a key enzyme in the synthesis of VLC-PUFAs, have been identified in Stargardt-like macular dystrophy (STGD3), a juvenile retinal dystrophy with macular deposits reminiscent of AMD^44–46^. Despite the biochemical, epidemiologic, and genetic evidence implicating PUFAs in AMD, the molecular mechanisms by which LC and VLC-PUFAs are involved in drusen formation, and AMD pathogenesis are still poorly understood. The finding that loss of ELOVL2 activity results in early accumulation of subRPE deposits strengthens the relationship between PUFAs and macular degeneration. Since *Elovl2* is expressed in both photoreceptors and RPE, whether these phenotypes of visual loss and subRPE deposits are due to cell autonomous function in the photoreceptors and RPE respectively or require interplay between photoreceptors and RPE still needs to be established.

## Conclusions

In summary, we have identified the lipid elongation enzyme ELOVL2 as a critical component in regulating molecular aging in the retina. Futher studies may lead to a better understanding of molecular mechanisms of aging in the eye, as well as lead to therapeutic strategies to treat a multitude of age-related eye diseases.

## ACKNOWLEDGEMENTS

We thank Dr. Trey Ideker for supporting work of T.W. We thank Ella Kothari and Jun Zhao in the UCSD Moores Cancer Center Transgenic Mouse Shared Resource for expert assistance in generation of edited mice. This work was supported by R01 EY02701 and RPB Special Scholar Award to D.S.K., by K12EY024225 to D.L.C. and by R01 GM086912 to B.A.H as well as by RPB Unrestricted Grant to Shiley Eye Institute. D.C., T. W., and K.D.R. were supported in part by a Ruth L. Kirschstein National Research Service Award (NRSA) Institutional Predoctoral Training Grant, T32 GM008666, from the National Institute of General Medical Sciences. Functional imaging and histology work were funded in part by the UCSD Vision Research Center Core Grant P30EY022589.

## METHODS

### Cell culture and treatment

WI38 and IMR-90 human fibroblasts were cultured in EMEM (ATCC) supplemented with 10% fetal bovine serum (Omega) and 1% penicillin/streptomycin (Gibco), and kept in a humidified incubator at 5% CO_2_ and 37°C. Confluence was calculated via ImageJ imaging software, including three fields of view per sample (10x). Upon confluence, cells were split and seeded at a 1:3 ratio. Population doublings (PD) were calculated by cell count. Knockdown lentivirus was generated using MISSION shRNA (Sigma) according to the manufacturer’s instructions. 5-Aza-2’-deoxycytidine was purchased from TSZ Chem (CAS#2353-33-5) and dissolved in cell culture medium at a concentration of 2μM. Cells were treated every day for a period of 48 hours. The medium was then replaced with regular cell culture medium, and the cells were cultured for 5 more days.

### Senescence-associated β-galactosidase (SA-β-gal) activity

The SA-β-gal activity in cultured cells was determined using the Senescence β-Galactosidase Staining Kit (Cell Signaling Technology), according to the manufacturer’s instructions. Cells were stained with DAPI afterward, and percentages of cells that stained positive were calculated with imaging software (Keyence), including three fields of view (10x).

### Nucleic acid analysis

DNA and RNA were isolated from human fibroblasts and mouse tissues with TRIzol (Ambion) according to the manufacturer’s instructions. RNA was converted to cDNA with iScript cDNA Synthesis Kit (Bio-Rad). qPCR was performed using SsoAdvanced Universal SYBR Green Supermix (Bio-Rad).

Methylated DNA Immunoprecipitation (MeDIP) was performed by shearing 1μg DNA by Bioruptor (Diagenode) for 8 cycles on the high setting, each cycle consisting of 30 seconds on and 30 seconds off. Sheared DNA was denatured, incubated with 1μg 5mC antibody MABE146 (Millipore) for 2 hours, then with SureBeads protein G beads (Bio-Rad) for 1 hour. After washing, DNA was purified with QIAquick PCR Purification Kit (Qiagen). qPCR was then performed as above. List of primers can be found in **Supplementary Table 1**

### Western Blotting

10μg of total protein isolated with TRIzol (Invitrogen) from retinas of WT mice of varying stages of development was subject to SDS-PAGE followed by Western blotting (see **Supp. Table 2** for antibodies used in the study). H3 served as loading control.

### Quantification of Western blots

WB ECL signals were imaged using BioRad ChemiDoc system. Background-subtracted signal intensities were calculated using ImageJ separately for ELOVL2 bands and H3 loading-control bands. ELOVL2 levels were calculated by dividing ELOVL2 signals by corresponding H3 signals, and then normalized to E15.5.

### RNAscope^®^ *In situ* hybridization

*In situ* hybridization was performed using the RNAscope^®^ Multiplex Fluorescent Assay v2 (ACD Diagnostics,Newark, CA). Mouse *Elovl2 Rpe65* and *Arr3* probes (p/n 542711, p/n 410151 and p/n 486551 respectively) were designed by the manufacturer. Briefly, fresh frozen histologic sections of mouse eyes were pretreated per manual using hydrogen peroxide and target retrieval reagents such as protease IV. Probes were then hybridized according to the protocol and then detected with TSA Plus®Fluorophores Fluorescein, Cyanine 3 and Cyanine 5 (Perkin Elmer, Waltham MA). Sections were mounted with DAPI and Prolong Gold antifade (ThermoFisher, Waltham, MA) with coverslip for imaging and imaged (Keyence BZ-X700).

### CRISPR-Cas9 design

CRISPR-Cas9 reagents were generated essentially as described^47^ and validated in our facility ^48^. T7 promoter was added to cloned Cas9 coding sequence by PCR amplification. The T7-Cas9 product was then gel purified and used as the template for in vitro transcription (IVT) using mMESSAGE mMACHINE T7 ULTRA kit (Life Technologies). T7 promoter and sgRNA sequence was synthesized as a long oligonucleotide (Ultramer, IDT) and amplified by PCR. The T7-sgRNA PCR product was gel purified and used as the template for IVT using the MEGAshortscript T7 kit (Life Technologies). A repair template encoding the C234W variant was synthesized as a single stranded oligonucleotide (Ultramer, IDT) and used without purification. Potential off-targets were identified using Cas-OFFinder^49^, selecting sites with fewest mismatches (http://www.rgenome.net/cas-offinder/). The founder mouse and all F1 mice were sequenced for off-targets. List of primers is in **Supplementary table 1**.

### Animal injection and analysis

All animal procedures were conducted with the approval of the Institutional Animal Care Committee at the University of California, San Diego (protocol number S17114). **CRISP/Cas9 injection**. C57BL/6N mouse zygotes were injected with CRISPR-Cas9 constructs. Oligos were injected into the cytoplasm of the zygotes at the pronuclei stage. Mice were housed on static racks in a conventional animal facility and were fed *ad libitum* with Teklad Global 2020X diet.

### Genotyping, mice substrains

To test for the potentially confounding Rd8 mutation, a mutation in the *Crb1* gene which can produce ocular disease phenotypes when homozygous, we sequenced all mice in our study for Rd8. C57BL/6J mice in the aging part of the study were purchased from the Jax laboratory and confirmed to be negative for mutation in *Crb1* gene. All C234W mutant animals and their littermates were heterozygous for Rd8 mutation. To test RPE65 gene, all animals were tested for the presence of the variants. All animals in the study harbor homozygous RPE65 variant Leu/Leu.

### Intravitreal injections

For the 5-Aza-dc injection study, mice were anesthetized by intraperitoneal injection of ketamine/xylazine (100 mg/kg and 10 mg/kg, respectively), and given an analgesic eye drop of Proparacaine (0.5%, Bausch & Lomb). Animals were intraocularly injected with 1μL of PBS in one eye, and 1 μL of 2μM 5-Aza-dc dissolved in PBS in the contralateral eye, every other week over a period of 3 months. Drug dosage was estimated based on our cell line experiments and on previously published data^50^.

**Autofluoresence imaging** was performed using the Spectralis ® HRA+OCT scanning laser ophthalmoscope (Heidelberg Engineering, (Franklin MA) as previously described (16) using blue light fluorescence feature (laser at 488 nm, barrier filter at 500 nm). Using a 55 degree lens, projection images of 10 frames per fundus were taken after centering around the optic nerve. The image that was most in focus was on the outer retina was then quantified blindly by two independent individuals.

**Electroretinograms (ERGs)** were performed following a previously reported protocol ^51^. Briefly, mice were dark-adapted for 12 h, anesthetized with a weight-based intraperitoneal injection of ketamine/xylazine, and given a dilating drop of Tropicamide (1.5%, Alcon) as well as a drop of Proparacaine (0.5%, Bausch & Lomb) as analgesic. Mice were examined with a full-field Ganzfeld bowl setup (Diagnosys LLC), with electrodes placed on each cornea, with a subcutaneous ground needle electrode placed in the tail, and a reference electrode in the mouth (Grass Telefactor, F-E2). Lubricant (Goniovisc 2.5%, HUB Pharmaceuticals) was used to provide contact of the electrodes with the eyes. Amplification (at 1–1,000 Hz bandpass, without notch filtering), stimuli presentation, and data acquisition are programmed and performed using the UTAS-E 3000 system (LKC Technologies). For scotopic ERG, the retina was stimulated with a xenon lamp at −2 and −0.5 log cd s/m2. For photopic ERG, mice were adapted to a background light of 1 log cd s/m2, and light stimulation was set at 1.5 log cd s/m2. Recordings were collected and averaged in manufacturer’s software (Veris, EDI) and processed in Excel.

### Immunostaining

Eyeballs were collected immediately after sacrificing mice, fixed in 4% paraformaldehyde for 2 hours, and stored in PBS at 4°C. For immunostainings, eyeballs were sectioned, mounted on slides, then incubated with 5% BSA 0.1% Triton-X PBS blocking solution for 1 hour. Primary antibodies (see **Supp. Table 2** for antibodies used in the study) were added 1:50 in 5%BSA PBS and incubated at 4°C for 16 hours. Following 3x PBS wash, secondary antibodies were added 1:1000 in 5%BSA PBS for 30 minutes at room temperature. Samples were then washed 3x with PBS, stained with DAPI for 5 minutes at room temperature, mounted, and imaged (Keyence BZ-X700).

### Lipid Analysis

Lipid extraction was performed by homogenization of tissues in a mixture of 1 mL PBS, 1 mL MeOH, and 2 mL CHCl3. Mixtures were vortexed and then centrifuged at 2200 g for 5 min to separate the aqueous and organic layer. The organic phase containing the extracted lipids was collected and dried under N2 and stored at −80 °C before LC-MS analysis. Extracted samples were dissolved in 100 μL CHCl3; 15 μL was injected for analysis. LC separation was achieved using a Bio-Bond 5U C4 column (Dikma). The LC solvents were as follows: buffer A, 95:5 water:methanol + 0.03% NH4OH; buffer B, 60:35:5 isopropanol:methanol: water + 0.03% NH4OH. A typical LC run consisted of the following for 70 minutes after injection: 0.1 mL/min 100% buffer A for 5 minutes, 0.4 mL/min linear gradient from 20% buffer B to 100% buffer B over 50 min, 0.5 mL/min 100% buffer B for 8 minutes and equilibration with 0.5 mL/min 100% buffer A for 7 minutes. FFA analysis was performed using a Thermo Scientific Q Exactive Plus fitted with a heated electrospray ionization source. The MS source parameters were 4kV spray voltage, with a probe temperature of 437.5°C and capillary temperature of 268.75°C. Full scan MS data was collected with a resolution of 70k, AGC target 1×106, max injection time of 100 ms and scan range 150–2000 m/z. Data-dependent MS (top 5 mode) was acquired with a resolution of 35 k, AGC target 1 × 105, max injection time of 50 ms, isolation window 1 m/z, scan range 200 to 2,000 m/z, stepped normalized collision energy (NCE) of 20, 30 and 40. Extracted ion chromatograms for each FFA was generated using a m/z ± 0.01 mass window around the calculated exact mass (i.e. palmitic acid, calculated exact mass for M-H is 255.2330 and the extracted ion chromatogram was 255.22–255.24). Quantification of the FFAs was performed by measuring the area under the peak and is reported as relative units (R.U.).

### Analysis of ELOVL2 promoter DNA methylation in mice and human

Reduced representation bisulfite sequencing (RRBS) in mouse blood was downloaded from Gene Expression Omnibus (GEO) using accession number GSE80672 ^52^. For each sample, reads obtained from sequencing were verified using FastQC ^53^, then trimmed 4bp using TrimGalore ^54^ (4bp) and aligned to a bisulfite-converted mouse genome (mm10, Ensembl) using Bismark (v0.14.3)^55^, which produced alignments with Bowtie2 (v2.1.0)^56^ with parameters “-score_min L,0,−0.2”. Methylation values for CpG sites were determined using MethylDackel (v0.2.1).

To explore methylation of the promoter region of ELOLV2, we first designated the promoter as −1000bp to +300bp with respect to the strand and transcription start site (TSS) and then identified profiled methylation CpGs using BEDtools (v2.25.0)^57^. We then binned each profiled CpG in the promoter region according to 30bp non-overlapping windows considering CpGs with at least 5 reads. We then grouped the 136 C57BL/6 control mice according to five quantile age bins, and took the average methylation for each age bin and each window. All analysis was performed using custom python (version 3.6) scripts, and plots were generated using matplotlib and seaborn.

To explore the homologous region in humans, we accessed human blood methylome data generated using the Human Illumina methylome array downloaded from GEO, using accessions GSE36054 ^58^ and GSE40279 ^2^ for a total of 736 samples. Methylation data were quantile normalized using Minfi^59^ and missing values were imputed using the Impute package in R. These values were adjusted for cell counts as previously described ^5^. To enable comparisons across different methylation array studies, we implemented beta-mixture quantile dilation (BMIQ)^5,60^ and used the median of the Hannum *et al*. dataset as the gold standard^2^.

We then identified probes within the promoter region of ELOLV2 in the human reference (hg19, UCSC), identifying 6 total probes in the commonly profiled region. We then grouped the 787 individuals according to 5 quantile age bins and grouped probes into 10bp non-overlapping windows. These data were then analyzed and plotted identically as for mice.

---

**Figure S1.**
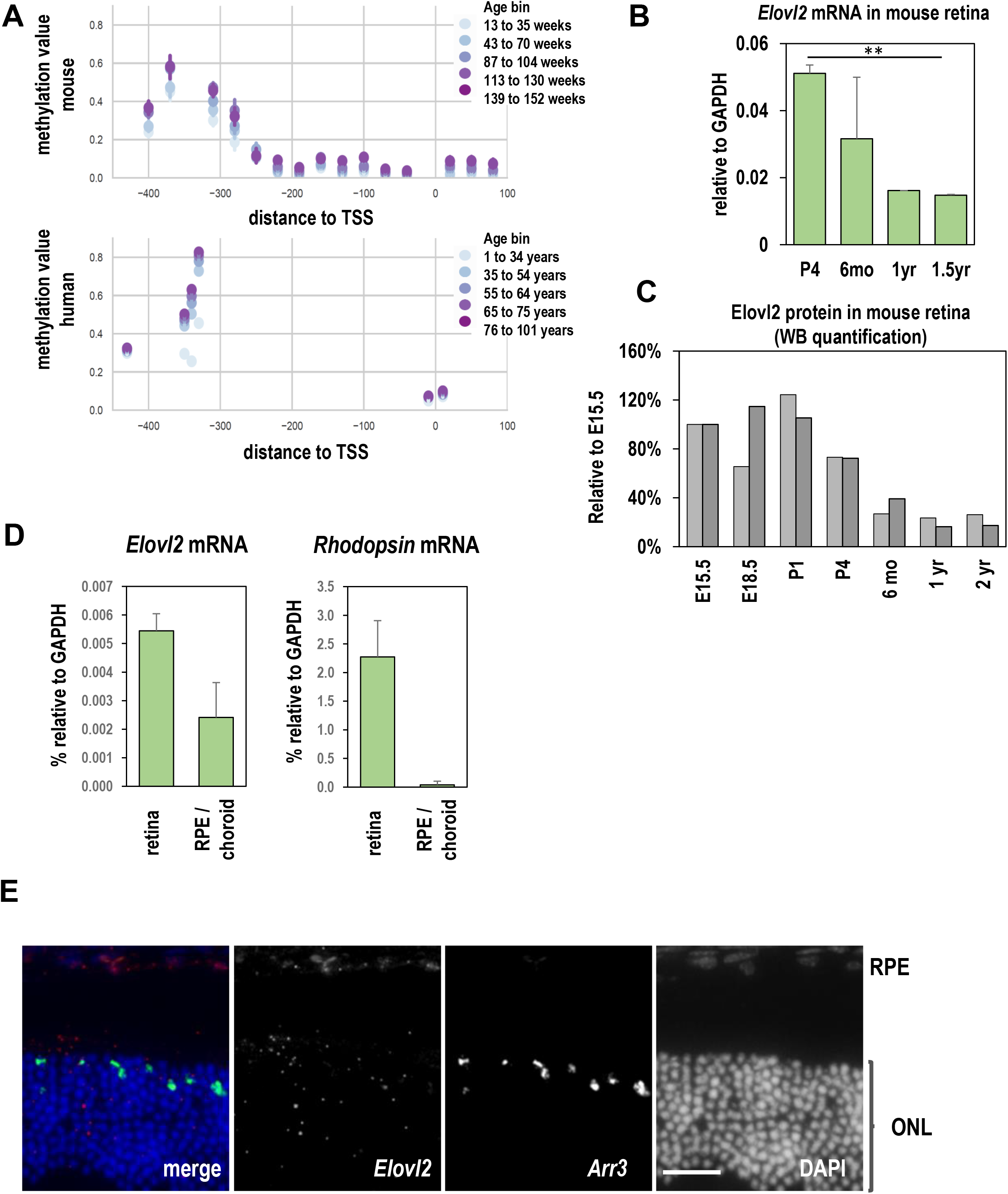
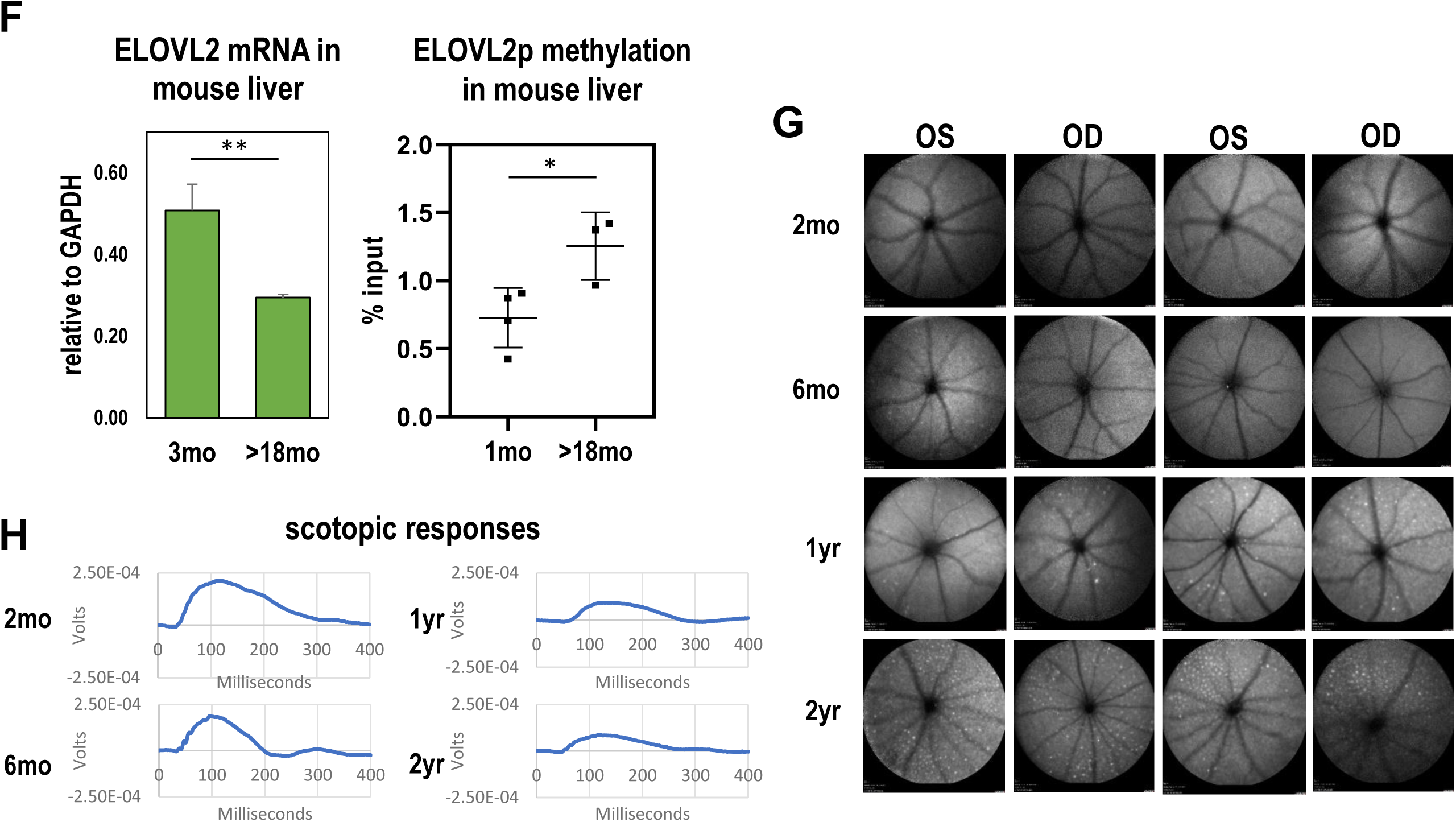
Aging characteristics in human and WT mice.

**A**. Top: ELOLV2 promoter methylation obtained using bisulfite-sequencing in mouse blood. The x-axis depicts the distance relative to the TSS of ELOLV2 (0) with respect to the direction of transcription according to 30bp non-overlapping windows. The y-axis depicts the methylation values obtained from 136 mice, which are grouped according to six quantile age bins, which are colored according to the legend by which darker colors reflect older age bins. The points reflect the average methylation value for each age bin and each window, and error bars represent the 95% confidence interval obtained from bootstrapping.

Bottom: The homologous region in human blood is depicted. These data were drawn from Illumina 450K human array platform where a total of 6 probes were identified in this region. To create an analogous representation, probes within 10bp of one another were averaged. 787 individuals were grouped according to five quantile age bins and the values are depicted in the same representation described above. **B**. Time course of Elovl*2* gene expression with age. N>=3, **p<0.01, 1-way ANOVA. **C**. Time course of retinal ELOVL2 protein expression by quantification of Western blots. Two identically performed experiments (Figure 1B) were quantified independently using ImageJ. **D**. *Elovl2* expression by qPCR in dissected mouse retina and RPE/choroid tissue. Rhodopsin, which is not expressed in RPE or choroid was used as purity control. N=3, error bars denote SD. **E**. RNAScope detecting expression of *Elovl2* and *Arr3* counterstained with DAPI in 3mo eyeballs show clear expression of *Elovl2* in RPE layer. Bar-50um. **F**. *Elovl2* expression and DNA methylation at *Elovl2* promoter in mouse liver young (3mo) and old mice (1.5-2y). N>=3, **p<0.01, *p<0.05, t-test. **G**. Autofluorescence images of WT mouse retinas at 2 months, 6 months, 1 year, and 2 years of age. **H**. Scotopic response of ERG in WT mice. Shown are representative traces obtained in retinas pictured in panel G.

**Figure S2.**
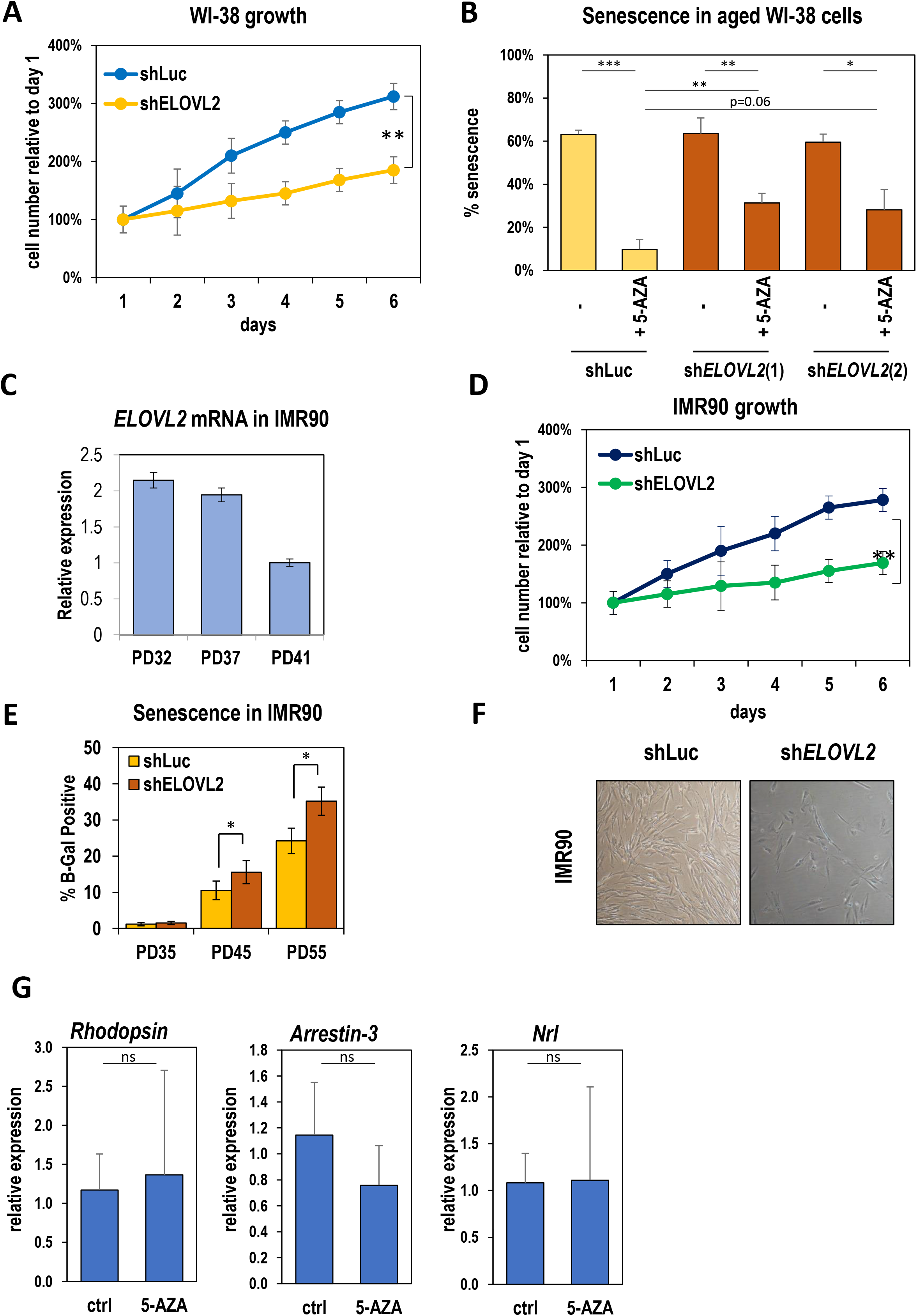
In vitro aging of WI38 and IMR90 cells.

**A**. Growth curves for WI38 cells upon either control knockdown (shLuc) or shRNA-mediated *ELOVL2* knockdown. N=3, ** p<0.01, t-test. B. Fraction of senescent WI-38 cells after addition with shLuc, shELOVL2(1) or shELOVL2(2) with 5-Aza. *, p<0.05 **p <0.01; ***p<0.001 **C**. *ELOVL2* expression by qPCR in IMR90 cells at PD32, 37 and 41. **D**. Growth curves in IMR90 cells, measured the same way as in panel A. **E**. Fraction of senescent cells measured by beta-galactosidase staining in IMR90 cells at given population doubling upon shRNA mediated knock-down of ELOVL2 gene or control Luc. Error bars denote SD, *, p<0.05. **F**. Morphology of in-vitro aging IMR90 cells upon either control knock-down (shLuc) or shRNA-mediated *ELOVL2* knockdown. **G**. *Rhodopsin, Arrestin-3 and Nrl* expression by qPCR after control (PBS) or 5-Aza-dc injection in 8mo mice. N=3, error bars denote SEM, ns, not significant.

**Figure S3.**
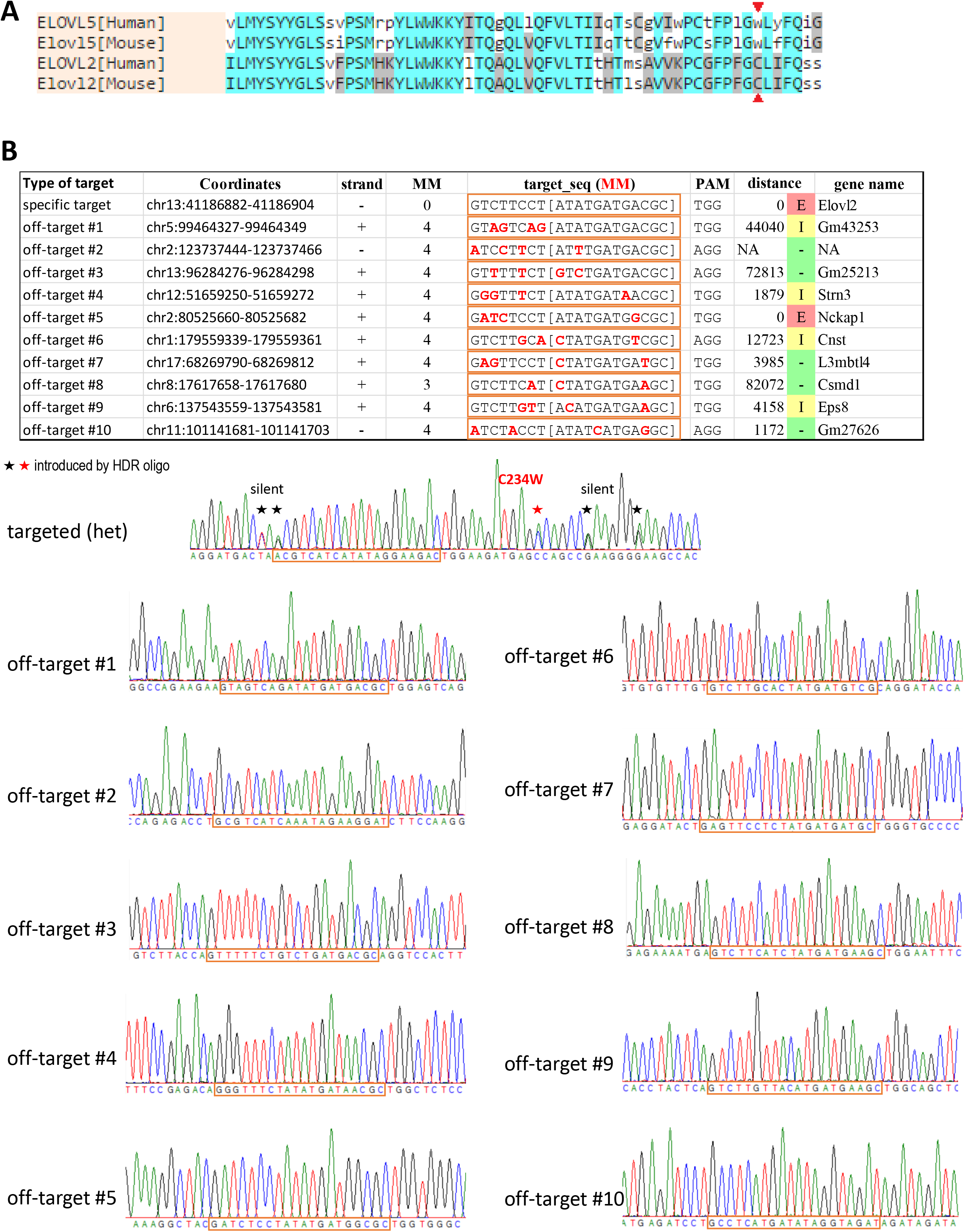
*Elovl2*^C234W^ mice.

**A**. ELOVL2 and ELOVL5 amino acid sequence similarity between human and mouse (aa 181-240). Red arrowheads denote targeted C234W mutation. **B**. Off-target analysis of *Elovl2* mutant mice. Specific and 10 top potential off-target sites are listed in a Table (top). Mismatches between the gRNA and target sites are shown in red. Bottom panels show sequencing chromatograms obtained from C234W +/-mice at the specific and potential off-target sites. Intended missense and silent substitutions are indicated by red and black asterisks, respectively. Off-target traces are identical to wild-type sequence indicating that no off-targeting occurred at these sites.

**Figure S4.**
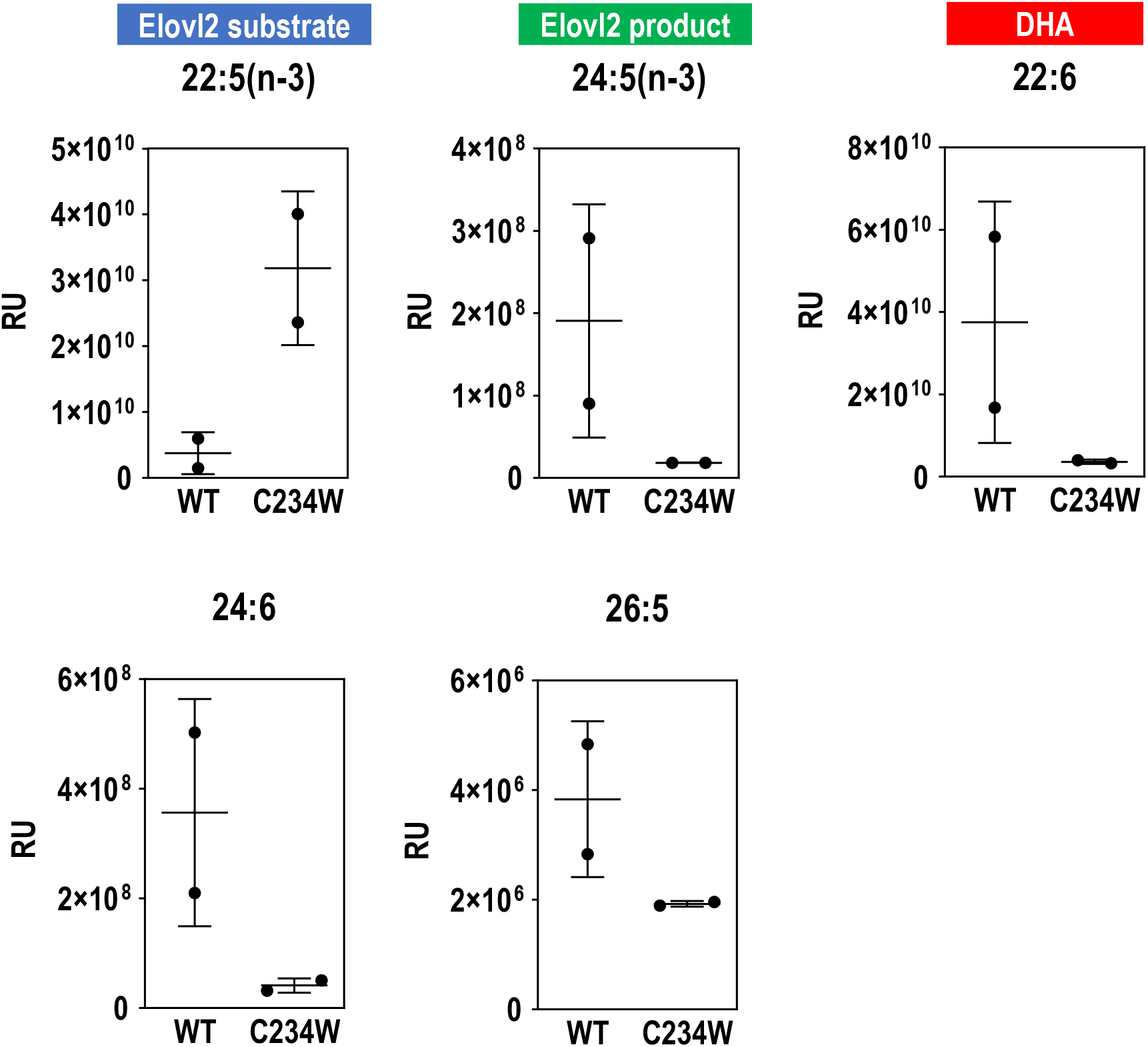
Lipid levels of ELOVL2 substrate (22:5(n-3)), product (24:5(n-3)) and downstream metabolites (22:6, a combination of DHA and 22:6(n-6), 24:6 and 26:5) measured in the liver of wild-type and ELOVL2 C234W mice. N=2. Error bars denote SD.

**Figure S5.**
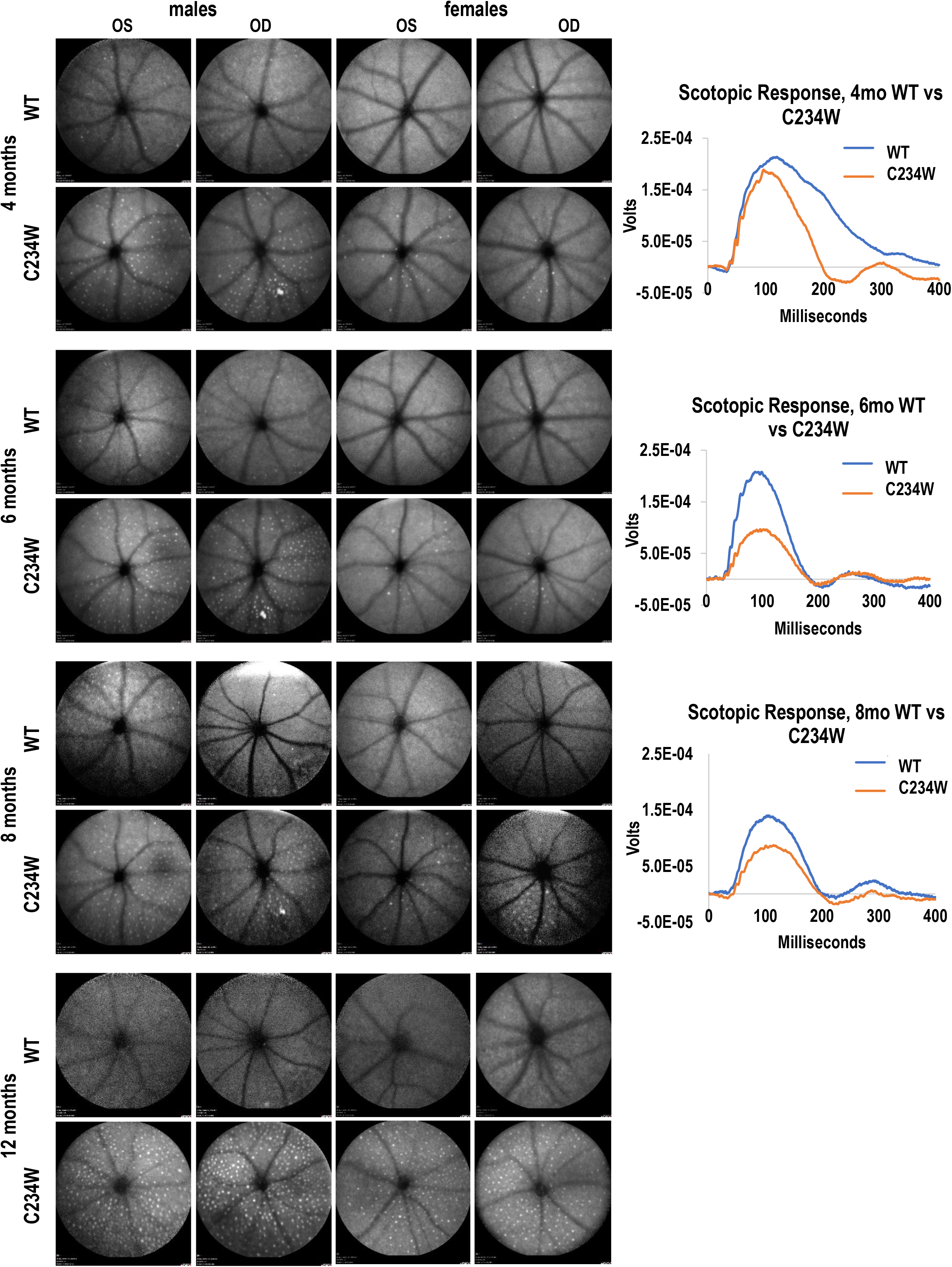
Fundus autofluorescence images of WT and *Elovl2*^C234W^ mice at 4, 6, 8 and 12 months with representative scotopic ERG waveforms (right panels). Number of autofluorescent deposits increases as animals age, what is accompanied by reduced maximal responses to visual stimuli.

**Figure S6.**
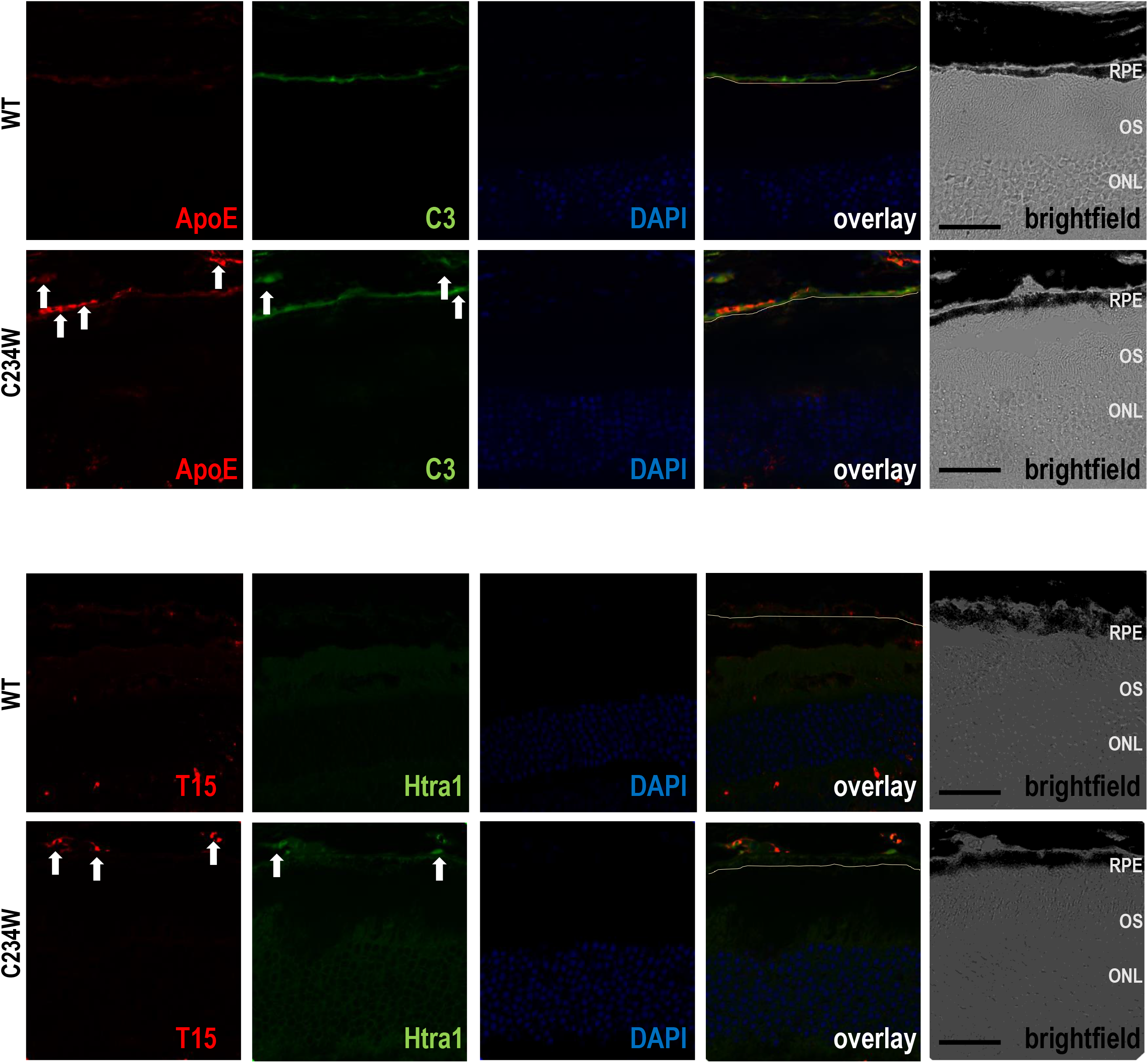
Characterization of sub-RPE aggregates by immunohistochemistry.

**A-D**. Immunostaining of Htra1, C3, ApoE, T-15 and C5b-9 counterstained with DAPI, in wild-type and C234W mouse retinas. Arrows indicate drusen-like aggregates. BF, bright-field, ONL, outer nuclear layer, INL, inner nuclear layer, RGC, retinal ganglion cells.

**Table S1.**
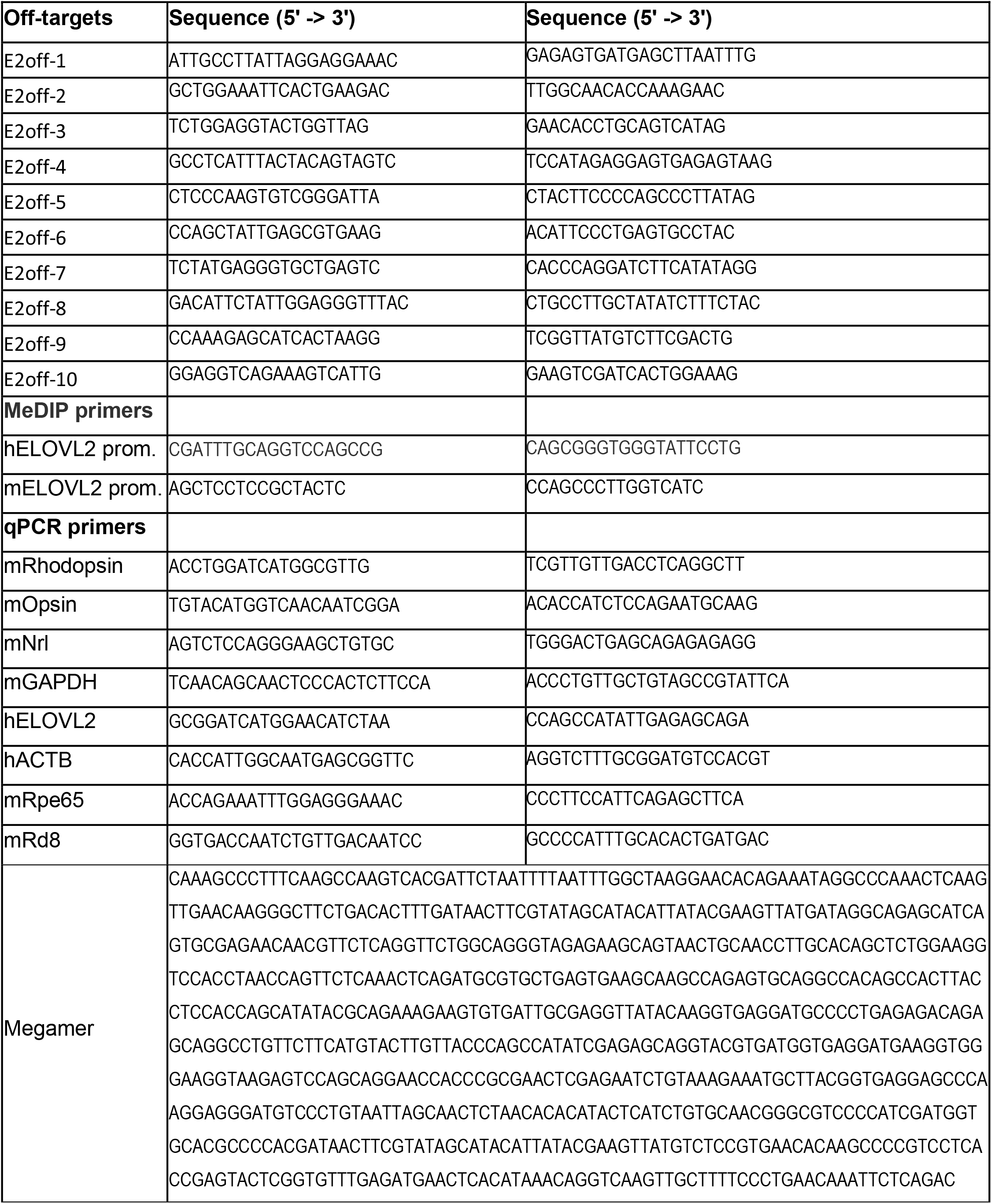

**Table S2.**

| **Immunostaining** | **Company, Cat#** | **RRID** |
| --- | --- | --- |
| TEPC 15 | Sigma M1421 | AB_1163630 |
| HtrA | Santa Cruz sc-377050 | AB_2813838 |
| C3 | Santa Cruz sc-58926 | AB_1119819 |
| C5-b9 | Santa Cruz sc-66190 | AB_1119840 |
| ApoE | Santa Cruz sc-13521 | AB_626691 |
| **MeDIP** |  |  |
| 5-methylcytosine | Millipore MABE146 | AB_10863148 |
| **Western blot** |  |  |
| ELOVL2 | Santa Cruz sc-54874 | AB_2262364 |
| Histone H3 | Cell Signaling 9715 | AB_331563 |

## Notes

#### Summary of Updates

AUTHOR LIST HAS BEEN UPDATED - Dorota Skowronska-Krawczyk should be the last and corresponding author.

